# TMS orientation and pulse waveform manipulation activates different neural populations: direct evidence from TMS-EEG

**DOI:** 10.1101/308981

**Authors:** Alberto Pisoni, Alessandra Vergallito, Giulia Mattavelli, Erica Varoli, Matteo Fecchio, Mario Rosanova, Adenauer G. Casali, Leonor J. Romero Lauro

## Abstract

Monophasic and biphasic TMS pulses and coil orientations produce different responses in terms of motor output and sensory perception. Those differences have been attributed to the activation of specific neural populations. However, up to date, direct evidence supporting this hypothesis is still missing since studies were mostly based on indirect measures of cortical activation, i.e., motor evoked potentials or phosphenes. Here, we investigated for the first time the impact of different coil orientations and waveforms on a non-primary cortical area, namely the premotor cortex, by measuring TMS evoked EEG potentials (TEPs). We aimed at determining whether TEPs produced by differently oriented biphasic and monophasic TMS pulses diverge and whether these differences are underpinned by the activation of specific neural populations. To do so, we applied TMS over the right premotor cortex with monophasic or biphasic waveforms oriented perpendicularly (in the anterior-posterior direction and vice-versa) or parallel (latero-medial or medio-laterally) to the target gyrus. EEG was concurrently recorded from 60 electrodes. We analyzed TEPs at the level of EEG sensors and cortical sources both in time and time-frequency domain. Biphasic pulses evoked larger early TEP components, which reflect cortical excitability properties of the underlying cortex, in both parallel directions when compared to the perpendicular conditions. Conversely, monophasic pulses, when oriented perpendicularly to the stimulated gyrus, elicited a greater N100, which is a reliable TEP component linked to GABAb-mediated inhibitory processes, than when parallel to the gyrus. Our results provide direct evidence supporting the hypothesis that TMS pulse waveform and TMS coil orientations affect which neural population is engaged.

## Introduction

Despite its large use for research and clinical purposes, different issues on the effects of Transcranial Magnetic Stimulation (TMS) on the cerebral cortex are still unknown (Triesch et al., 2015). One interesting methodological aspect concerns the differences in neurophysiological responses evoked by monophasic and biphasic stimulation pulses. In the monophasic mode, a strong initial current flow is followed by a smaller current in the opposite direction. The initial flow has a quick peak (about 50 μs after pulse onset) and effectively excites neurons, while the subsequent return current, which lasts several hundreds of μs, does not elicit action potentials (Groppa et al., 2012). The biphasic pulse, instead, has a cosine waveform: an initial peak is followed by a reversal current and by another subsequent peak. In this pulse configuration, each phase of the pulse induces an effective stimulation, which spreads in the same or opposite direction as the initial one. In the biphasic mode, then, all pulse phases are effective in stimulating the cortical nervous tissue but seem to involve different neuronal populations (Groppa et al., 2012), even if the second half cycle is more effective.

It has been reported that these two TMS pulse waveforms induce differential electrophysiological outputs as measured by Motor Evoked Potentials (MEPs; e.g., Kammer et al., 2001; Niehaus et al., 2000; Sommer et al., 2006) and can induce distinct plastic after-effects following repetitive TMS protocols (Pascual-Leone et al., 1994; Sommer et al., 2002; Tings et al., 2005). The comparison between single monophasic and biphasic pulses over the motor cortex (M1) suggests that at a given amplitude of the initial current, biphasic stimulation is more effective than the monophasic one in eliciting MEPs (Kammer et al., 2001). For rTMS, the effect of the two pulse configurations seems to be reversed. Reports, indeed, showed that the inhibition of M1 excitability exerted by monophasic low-frequency rTMS (1HZ-rTMS) is more prolonged compared to the biphasic one (Sommer et al., 2002; Taylor & Loo, 2007), especially when the current is oriented in the anterior-posterior direction (Tings et al., 2005). Conversely, monophasic posterior-anterior rTMS induces an increase of M1 cortical excitability as, to a lesser extent, also did the latero-medial orientation (Tings et al., 2005). Similar results were also obtained over primary visual cortex (V1), where low-frequency monophasic rTMS decreased contrast sensitivity for visual stimuli after stimulation (Antal et al., 2002). Some authors suggested that monophasic rTMS activates a relatively uniform neural population and could therefore be more effective in producing sustained plastic after-effects. Conversely, biphasic pulses generate a more complex pattern of neural activations (Arai et al., 2005, 2007; Sommer et al., 2002), reducing the overall stimulation-induced after-effects. The ultimate reason for these differences, however, is still debated (Sommer et al., 2006) and so is the role of the different components of biphasic pulses for in vivo and in vitro studies (Kammer et al., 2001; Maccabee et al., 1998). Another compelling issue is the orientation of the TMS-induced electric field, which indeed has been associated with different neurophysiological responses, due to dissimilar activations of the underlying neural population or by recruiting different neurons (for a review see Di Lazzaro et al., 2008).

Crucially, research focused on the neurophysiological changes occurring in M1, since motor cortex activation can be indirectly recorded via MEPs or cortico-spinal recordings, or in V1, by analyzing phosphenes perception. However, TMS protocols are widely used in research (see Luber & Lisanby, 2014 for a review) and clinical protocols (see Lefaucheur et al., 2014 for a review) in a variety of areas, which have very different cytoarchitectonic and neurophysiological properties and cannot be directly compared with M1 or V1 (Taylor & Loo, 2007). A further element preventing a clear understanding of the processes underlying different TMS protocols is that studies often report coil orientation referred to the subjects’ head, only inferring the underlying cortical geometry. For example, the 45° angle usually applied for an optimal M1 stimulation, is reported to be perpendicular to the precentral gyrus, but a vast number of studies do not have individual MRIs to check if this is true (Sparing et al., 2010).

Improving our knowledge of TMS mechanisms can be useful to optimize research and treatment protocols. In this perspective, we used a navigated TMS combined with electroencephalography (TMS-EEG) system to investigate the impact of TMS pulse waveform and coil orientation on cortical excitability. Specifically, we compared monophasic and biphasic pulses delivered over the right premotor cortex orienting the coil perpendicularly (i.e. applying an anterior-posterior pulse, A – P, or posterior-anterior one, P – A) or parallel (i.e. applying a latero-medial, L – M, or medio-lateral, M – L, pulse) to the right premotor cortex, localized on individual MRIs. The main advantage of TMS-EEG approach is to provide real-time and direct information on cortical reactivity through TMS-evoked potentials (TEPs) recording. TEPs have been indeed consistently reported as being an informative measure of cortical excitability (Casarotto et al., 2010; Ilmoniemi et al., 1997; Ilmoniemi & Kičić, 2010; Lioumis et al., 2009) and connectivity (Komssi et al., 2002, 2004, 2007; Massimini et al., 2005; Mattavelli et al., 2013, 2016; Pisoni et al., 2017; Romero Lauro et al., 2014, 2016). Interestingly, previous research linked TEPs components with specific neurophysiological properties of the stimulated cortical area. In particular, early TEP components (0-60ms) have been described as reflecting different processes: on one hand, ipsilateral components reflected excitability properties of the targeted region, being affected by neurophysiological interventions aimed at modulating synaptic strength and cortical excitability such as tDCS (Pellicciari et al., 2013; Pisoni et al., 2017) or rTMS (Esser et al., 2006; Veniero et al., 2010, 2012); conversely, a component peaking at ~45ms and centered on contralateral homologue areas seems to be involved in GABAa mediated intracortical inhibition (Premoli et al., 2014; Ziemann et al., 2015). Expressly for the premotor cortex, one of the most consistently described components peaks around 100 ms from TMS pulse, and is related to GABAb cortico-cortical inhibition (Ilmoniemi & Kičić, 2010; Kičić et al., 2008; Premoli et al., 2014). By systematically varying TMS pulse waveform, orientation, and direction we thus aimed at investigating possible differences in TEPs components and amplitude to assess which neurophysiological responses are triggered by different stimulation protocols and to measure their efficacy in eliciting cortical responses.

## Material and ethods

### Participants

Ten healthy, right-handed volunteers (6 male, mean age 28.3, SD 6.4, range 21-39) participated in the study. Each participant completed an Adult Safety Screening Questionnaire (Keel et al., 2001) and gave informed written consent before study procedures. Participants with any contraindication, such as brain injury or surgery, heart attack or stroke and use of medications known to alter cortical excitability (e.g., anti-depressant medication), were excluded (Rossi et al., 2009). Each subject had an individual structural MRI of the brain to be used for neuronavigation. The study was performed in the TMS-EEG laboratory of the University of Milano-Bicocca and was approved by the local Ethic Committee.

### Procedure

Each participant took part in two experimental sessions separated by ~6 months between them, each consisting of four TMS-EEG recordings performed with different TMS parameters, all delivered over the right premotor cortex (Fig.1a), varying for both TMS waveform (monophasic vs biphasic), orientation over the target gyrus (parallel vs perpendicular), and direction (A – P and P – A for the perpendicular orientation and M – L and L – M for the parallel orientation). Each recording lasted about seven mins, during which participants fixated a white cross in black screen (17”) in front of them. Around 180 TEPs were recorded in each condition that, as previously reported, allows having a good number of artefact-free trials for analysis (Casarotto et al., 2010; Pisoni et al., 2017; Romero Lauro et al., 2014). The order of the eight TMS conditions was quasi counterbalanced across subjects, meaning that waveforms and orientations were counterbalanced, while the A – P and L – M directions were always performed in session 1 and P – A and M – L directions in session 2.

**Fig. 1:**
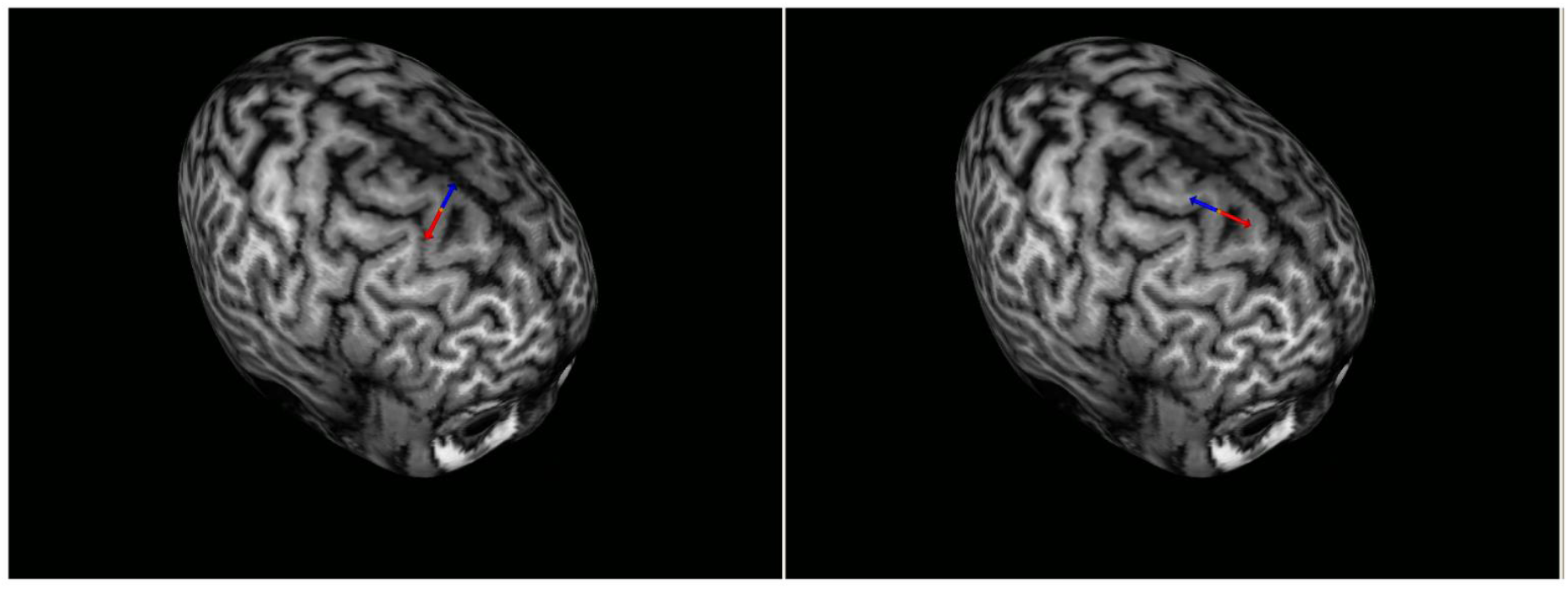
NBS screenshots displaying stimulation position and direction in the parallel (M – L, left) and perpendicular (P – A, right) orientations. The red arrow indicates the direction of the strongest phase of the stimulation (i.e., the only one in the monophasic and the second one of the biphasic pulse).

### TMS stimulation

TMS was delivered using an Eximia^TM^ TMS stimulator (Nexstim^TM^, Helsinki, Finland) using one monophasic and one biphasic focal figure of eight 70-mm coils. The stimulation target was the right premotor area. High-resolution (1×1×1 mm) structural magnetic resonance images (MRI) were acquired for each participant using a 3 T Intera Philips body scanner (Philips Medical Systems, Best, NL). TMS target was identified on individual MRIs using an integrated Navigated Brain Stimulation (NBS) system (Nexstim^TM^, Helsinki, Finland) which employs infrared-based frameless stereotaxy, to map the position of the coil and the participant’s head, within the reference space of the individual’s MRI space. The NBS system allowed to continuously monitor the position and orientation of the coil, thus assuring precision and reproducibility of the stimulation across recordings. The importance of using NBS with individualized MRIs rather than adjusting the coil angle and position according to scalp landmarks is crucial to correctly target the desired cortical area, which may be differently displaced in each subject (Sparing et al., 2010). No individual functional MRI was acquired, thus the functional specificity of the stimulation area could not be assessed. The NBS system estimated online the intensity (V/m) of the intracranial electric field induced by TMS at the stimulation hotspot, accounting for the head and brain shape of each participant, and taking into consideration the distance from scalp and coil position. Resting Motor Threshold, indeed, could have been a misleading measure to calibrate stimulation intensity, since the cortical thickness and cortical reactivity may greatly vary between M1 and the premotor cortex (e.g., see Kähkönen et al., 2005; Peterchev et al., 2012). We thus calibrated TMS intensity considering the estimated induced cortical electric field and checking, in a short preliminary recording before each real session, whether an on-line response of at least ~ 2 μV could be evoked, by starting at an estimated intensity of the electrical induced field of 90 V/m. Mean estimated electric field at the stimulation target for all condition are reported in Tab. 3. Critically, mean estimated induced electric field did not differ between any of the stimulation protocols (all ps>.14). The corresponding mean stimulation intensities, expressed as a percentage of the maximal output of the stimulator, are reported in Tab. 4. Crucially, within monophasic and biphasic conditions, mean stimulator output did not differ between the four different directions (all ps>=.08). The coil was tangentially placed to the scalp, and adjusted for each participant to direct the electric field parallel (L – M or M – L) or perpendicular (A – P or P – A) to the shape of the cortical gyrus (See Fig.1). The stimulation direction is relative to the first and unique cycle of the monophasic and the second, strongest cycle of the biphasic pulse. As in previous studies (Julkunen et al., 2008; Mütanen et al., 2013; Zanon et al., 2013), the stimulation of the premotor cortex did not evoke evident facial muscular artefacts, and no twitch of the contralateral upper limbs were indeed reported for any subject. TMS pulses were delivered at an inter-stimulus interval randomly jittering between 2000 and 2300 ms.

### EEG Recording during TMS

EEG signal was continuously recorded using a TMS compatible 60-channels amplifier (Nexstim Ltd., Helsinki, Finland), which prevents saturation using a proprietary sample-and-hold circuit which holds the amplifier output constant from 100 μs pre to 2 ms post-TMS pulse (Virtanen et al., 1999). Two extra-electrodes placed over the forehead were used as ground. Eye movements were recorded using two additional electrodes placed near the eyes to monitor ocular artefacts. As in previous studies, during EEG recordings, participants wore earplugs and heard a continuous masking noise to cover TMS coil discharge (Casarotto et al., 2010; Massimini et al., 2005; Romero Lauro et al., 2014), avoiding thus the emergence of auditory evoked potentials (Ter Braack et al., 2015). Electrodes impedance was kept below five kΩ, and EEG signals were recorded with a sampling rate of 1450 Hz and in common reference.

Data pre-processing was carried out using Matlab R2012a (Mathworks, Natick, MA, USA). EEG recordings were band-pass filtered between 1 and 45 Hz. Then, EEG signals were split into epochs starting 800 ms before and ending 800 ms after the pulse. EEG signals were down-sampled to 725 Hz. Trials with excessive artefacts were removed by a semi-automatic procedure and visual inspection (Casali et al., 2010) and TEPs were computed by averaging selected artefact-free single epochs. Bad channels were interpolated using spherical interpolation function of EEGLAB (Delorme & Makeig, 2004). TEPs were then averaged-referenced and baseline corrected between −300 and −50 ms before the TMS pulse. Independent component analysis (ICA) was applied to remove residual artefacts (Delorme et al., 2007). The average number of accepted trials considered in the analysis is reported in supplementary Tab. 1 of the Supplementary Materials while the mean signal to noise ratio, which was greater than 1.55 in all sessions (Casali et al., 2013), is reported in Tab. 2 of the Supplementary Materials.

To assess where and when cortical responses to TMS differed within pulse waveform according to coil orientation, TEPs were rectified and compared through a cluster-based test (Maris & Oostenveld, 2007) as implemented in the FieldTrip MATLAB toolbox for M/EEG analysis (freely available at http://fieldtrip.fcdonders.nl/; Oostenveld et al., 2011). Specifically, a whole-head, cluster-based permutation-corrected t-test was run between A – P, P – A, M – L, and L – M orientations within each coil type. To assess differences in amplitude, this procedure performs 10000 permutations by shuffling trial labels. Then, for each permutation, independent sample t-tests are performed at each time-point. All samples with a statistic corresponding to a P-value smaller than.05 are clustered together by spatial proximity. Finally, the cluster statistic is computed by taking the sum of the t-values within each cluster. The cluster-corrected threshold is then obtained by computing the permutation distribution of the maximum cluster statistic. This procedure thus corrects for multiple comparisons by permuting the data and clustering them based on their spatial and temporal proximity. In our case, for each comparison, permutations were performed for the whole 0-250ms time window with a permutation-significant level of p=.05. Critically six comparisons were performed within each waveform: A – P vs. P – A; M – L vs. L – M; A – P vs. M – L; A – P vs. L – M; P – A vs. L – M; P – A vs. M – L.

To better refine where the cortical activation induced by the different TMS protocols was taking place, source analysis was performed. Firstly, individual standardized meshes were reconstructed for each participant, starting from their structural MRIs (SPM8, Ashburner et al., 2011; Litvak et al., 2011), obtaining meshes of cortex, skull and scalp compartments (containing 3004, 2000 and 2000 vertices, respectively), normalized to the Montreal Neurological Institute (MNI) atlas (Casali et al., 2010). Then, for each participant, EEG sensor position was aligned to the canonical anatomical markers (pre-auricular points and nasion), and the forward model was computed. The inverse solution was computed on the average of all artefact-free TMS-EEG trials using the weighted minimum norm estimate with smoothness prior, following the same procedures as in Casali et al. (2010). This method is advantageous because it provides stable solution also in the presence of noise (Silva et al., 2004), and does not require any a priori assumption about the nature of the source distribution (Hämäläinen & Ilmoniemi, 1994). After source reconstruction, a statistical threshold was computed to assess when and where the post-TMS cortical response differed from pre-TMS activity (i.e., to identify TMS-evoked response). To do so, a nonparametric permutation-based procedure was applied (Pantazis et al., 2003). Binary spatial-temporal distribution of statistically significant sources was obtained, and thus only information from significant cortical sources was used for further analyses. As a measure of global cortical activation, we cumulated the absolute Significant Current Density (global SCD, measured in mA/mm2, Casali et al., 2010) over all 3004 cortical vertexes and over three time windows encompassing the TEPs components under investigation (0-30 ms, 30-80 ms and 80-130 ms) for each TMS protocol. Moreover, a local SCD in the vertexes within the BA targeted by TMS, i.e., right BA 6 identified using an automatic tool of anatomical classification (WFUPickAtlas tool; http://www.ansir.wfubmc.edu), was computed. Finally, an index of current scattering (SCS, Casali et al., 2010) was computed to estimate how spread the induced signal was. These measures were compared between stimulation protocols, within each coil type, using a series of linear mixed models (Baayen et al., 2008) in R computing environment (R Development Core Team, 2008). Indices derived from the source modeling were included in the model as dependent variables, while the stimulation protocol inclusion as fixed effect was estimated with an LRT procedure (Baayen et al., 2008), including it in the final model only if it significantly increased the model’s goodness of fit. The by-subject random slope was included as random factor.

Finally, to investigate whether the TMS evoked responses differed not only in amplitude, but also in spectral components, a time-frequency analysis was run. In particular, within Fieldtrip toolbox, the TMS induced time-frequency representations (TFR) of power was computed by convolving single trials with Morlet wavelets that had a width of 3.5 cycles (see Rosanova et al., 2009 for a similar procedure). Wavelet convolution was done between 2 and 40Hz, in 2Hz steps, and a time step of 2ms between −800ms and +800ms around the TMS pulse. To assess the significance of differences in oscillation power across conditions, the same procedure adopted for the source analyses data was used. In particular, mean power values mediated from the 4 electrodes near the TMS coil (Fz, F2, FCz, FC2) were computed for 4 different frequency bands (θ: 4-7 Hz; α: 8-12 Hz; β: 13:30 Hz; high β: 31-40 Hz), were baseline normalized, and finally were cumulated over the whole TEP time window (0-250ms). These values were used for the linear mixed regression previously described, comparing the four monophasic and the four biphasic directions separately.

## Results

### Effects of coil orientation on TEPs evoked by monophasic pulses

#### Amplitude at the scalp level

To specifically assess which part of the evoked cortical response differed according to coil orientation, the amplitude of the evoked response was compared with a cluster based permutation t-test on the whole 0-250ms TEP duration.

Fig. 2 shows the unrectified grand average of TEPs in the four monophasic conditions. As the first components, with a latency comprised within the first 70ms, have a similar time-course and amplitude in all orientations and directions, responses start to differ in the component peaking around 100ms, where the perpendicular orientations, in both A – P and P – A directions, elicit a greater response as compared to the parallel orientations, similarly in the M – L and L – M directions. The following late component, peaking around 200 ms, seems greater for the A-P orientation, even if a consistent response is also recorded for the L-M direction.

**Fig. 2:**
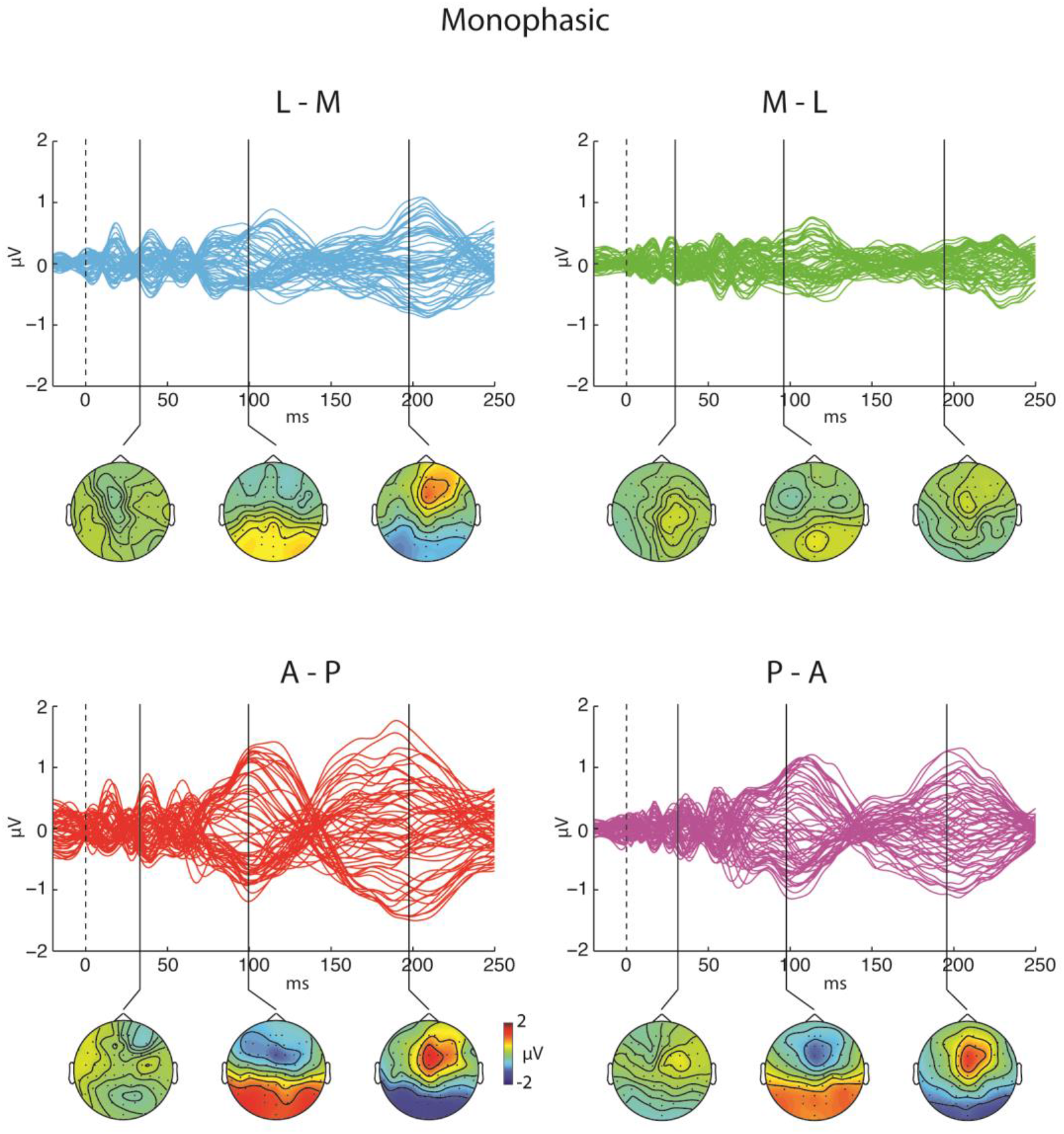
Butterfly plots and scalp topographies of TEPs grand average triggered by monophasic TMS. Top line shows results for the L – M and M – L directions while bottom line shows results for the A – P and P – A directions.

The cluster-based analysis confirmed these observations. No significant differences were present for any components between the two directions within parallel and perpendicular orientations. By contrast, when comparing the two orientations, A – P stimulation resulted in a greater 100ms component when delivering L – M (significant cluster from 80 to 130ms, p<.001) and M – L monophasic TMS (significant cluster from 50 to 160ms, p<.001). Moreover, this latter comparison showed that A – P stimulation triggered greater components in a significant early cluster between 170 and 240ms (p=.014) and an earlier one, between 20 and 40ms (p=.038). The topography of these differences, centred around 100ms post TMS (Fig. 3), shows higher evoked activity in the perpendicular A – P direction over a large cluster of electrodes, comprising a set of sensors located in the fronto-central areas, including the region under the TMS coil and its contralateral homologue when compared to both the parallel L – M and M – L directions.

**Fig. 3:**
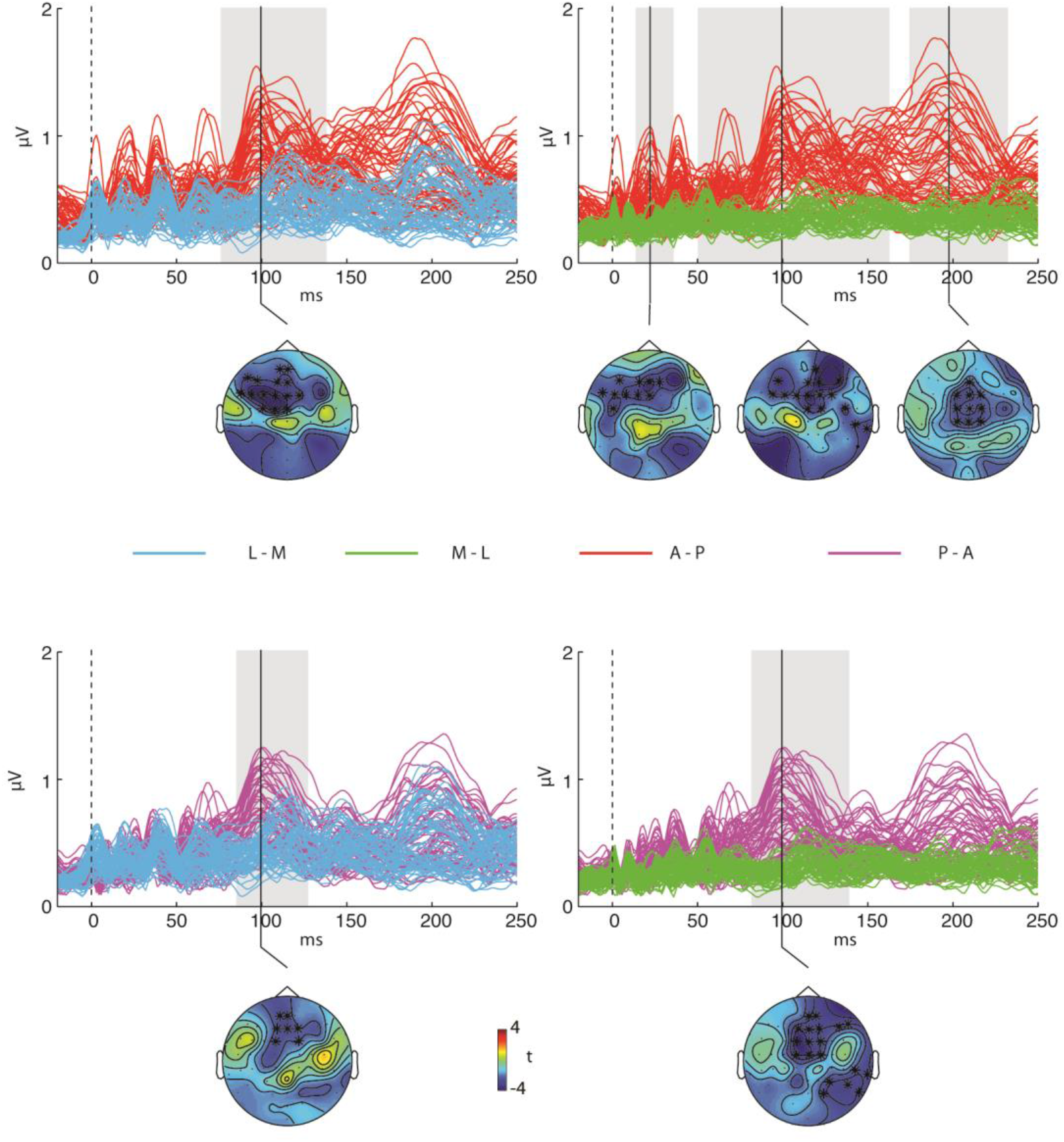
Results of the cluster based analysis for the monophasic pulse waveform. Top line displays a superimposition of rectified TEPs for the L – M and A – P (left) and M – L and A – P conditions (right). Grey shaded areas represent time windows in which the two waveforms statistically differ. Scalp topographies represent the distribution of the statistics for each comparison with significant electrodes plotted in bold. The same is reported in the bottom line for the comparison between P – A and L – M (left) and M-L (right) protocols.

Similarly, P – A monophasic TMS elicited greater 100ms component compared to both L – M (significant cluster 90-110ms, p=.05) and M – L (significant cluster 80-130ms, p=.006) directions (see Fig. 3).

#### Cortical Source modeling

Confirming the results at the sensors level, global SCD was smaller in the late component (80-130ms) for the L – M protocol compared to both A – P (χ^2^ (1)= 6.61; p=.01) and P – A (χ^2^ (1)= 5.25; p=.02) perpendicular orientations. M – L protocol, instead, showed marginal differences with the A – P and P – A directions (χ^2^ (1)= 3.4; p=.06 and χ^2^ (1)= 3.3; p=.06 respectively; see Fig. 4). Local SCD, computed in BA6, which was targeted by TMS, showed a clear increase in the induced current in the 100ms component for the perpendicular orientations compared to the parallel ones. In particular, A – P protocol elicited a greater late component compared to both L – M (χ^2^ (1)= 4.81; p=.028) and M – L (χ^2^ (1)= 5.49; p=.019) parallel directions. Similarly, the P – A stimulation induced a greater 100ms component compared to the L – M (χ^2^ (1)= 5.44; p=.02) and M – L (χ^2^ (1)= 7.63; p=.005) protocols (See Fig. 4). No other significant difference was present for the early components.

**Fig. 4:**
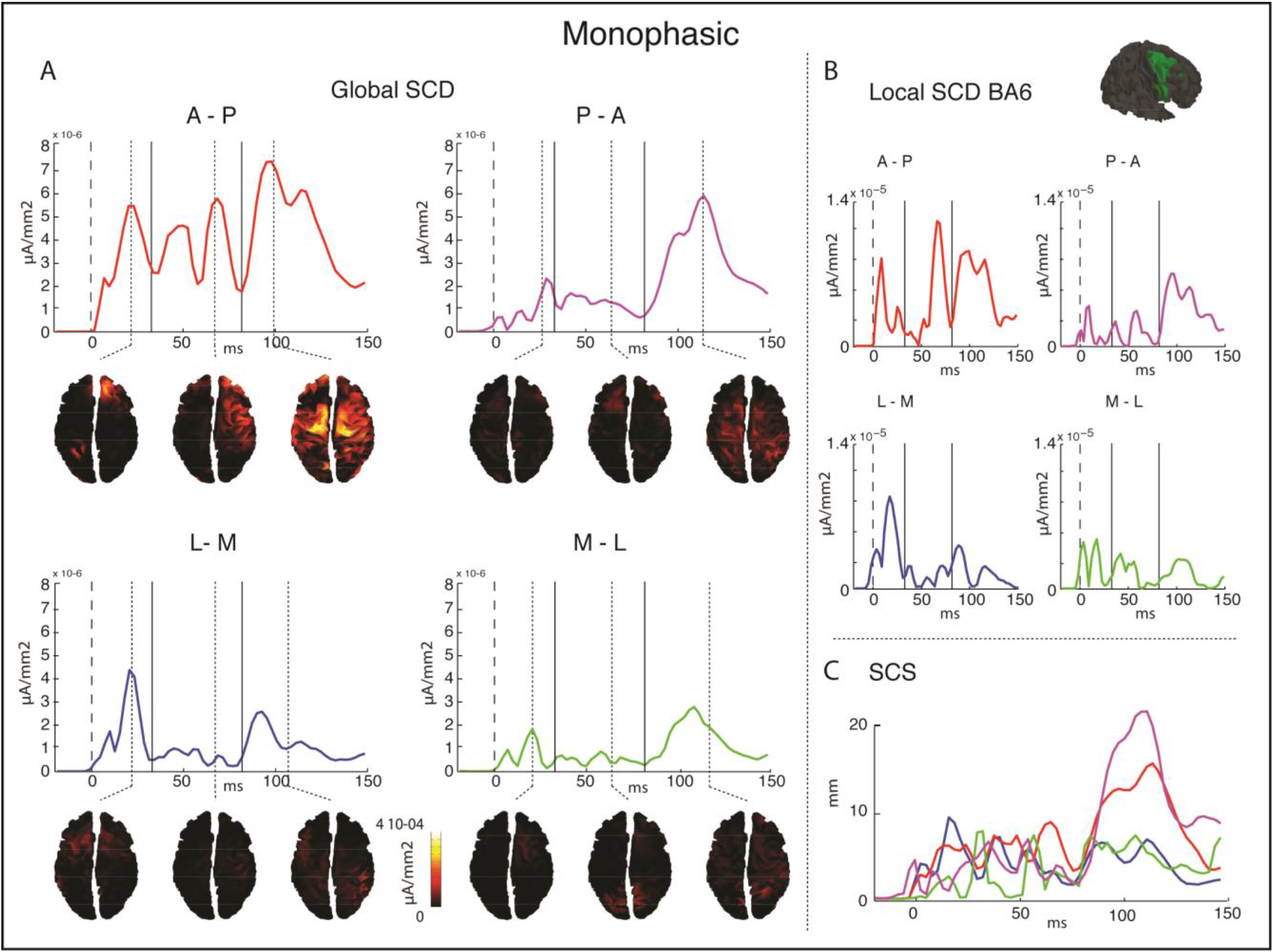
Results of the source analyses for the monophasic protocols. A) plot of the global SCD for the four directions with cortical source reconstruction at the local maxima for each of the three analyzed time windows (0-30ms; 30-80ms; 80-150ms). B) Plot of local SCD in the right BA6. C) plot of the SCS for the 4 protocols.

The Significant Current Scattering confirmed that, while in the first and second time-windows no difference in the spread of the induced current was present between the four protocols, in the third time window current spreads more over the cortex in the perpendicular compared to the parallel orientations. In particular, in the A – P direction SCS was greater compared to both L – M (χ^2^ (1)= 5.6;p=.017) and M – L (χ^2^ (1)= 3.9; p=.049) conditions. Similarly, the P – A protocol induced more spread cortical activity compared to the L – M condition (χ^2^ (1)=5; p=.025) and marginally compared to the M – L one (χ^2^ (1)= 3.6, p=.056). In the first time-window, instead, the A – P protocol induced more widespread activity compared to the P – A direction.

#### Time-Frequency analysis

Time-Frequency analysis showed a different effect of TMS orientation and direction on the oscillatory cortical response. A – P stimulation induced greater oscillation in the theta, alpha, beta and high beta bands compared to both L – M (θ: p<.001; α: p<.001; β: p<.001; high β: p=.013) and M – L (θ: p=.02; α: p=.046; β: p=.018 high β: p<.001) parallel directions. Conversely, P – A perpendicular stimulation elicited greater activity in all the considered frequency bands compared only to the L – M direction (θ: p=.029; α: p=.046; β: p=.05). Perpendicular stimulation differed in the alpha, beta and high beta bands, with A – P direction eliciting greater power in both frequency bands compared to P – A direction (α: p=.024; β: p=.001; high β: p=.015). Conversely, parallel directions differed in the alpha (p=.025) and beta (p=.007) bands, with a higher power in the L – M compared to the M – L direction (see Fig 5).

**Fig 5:**
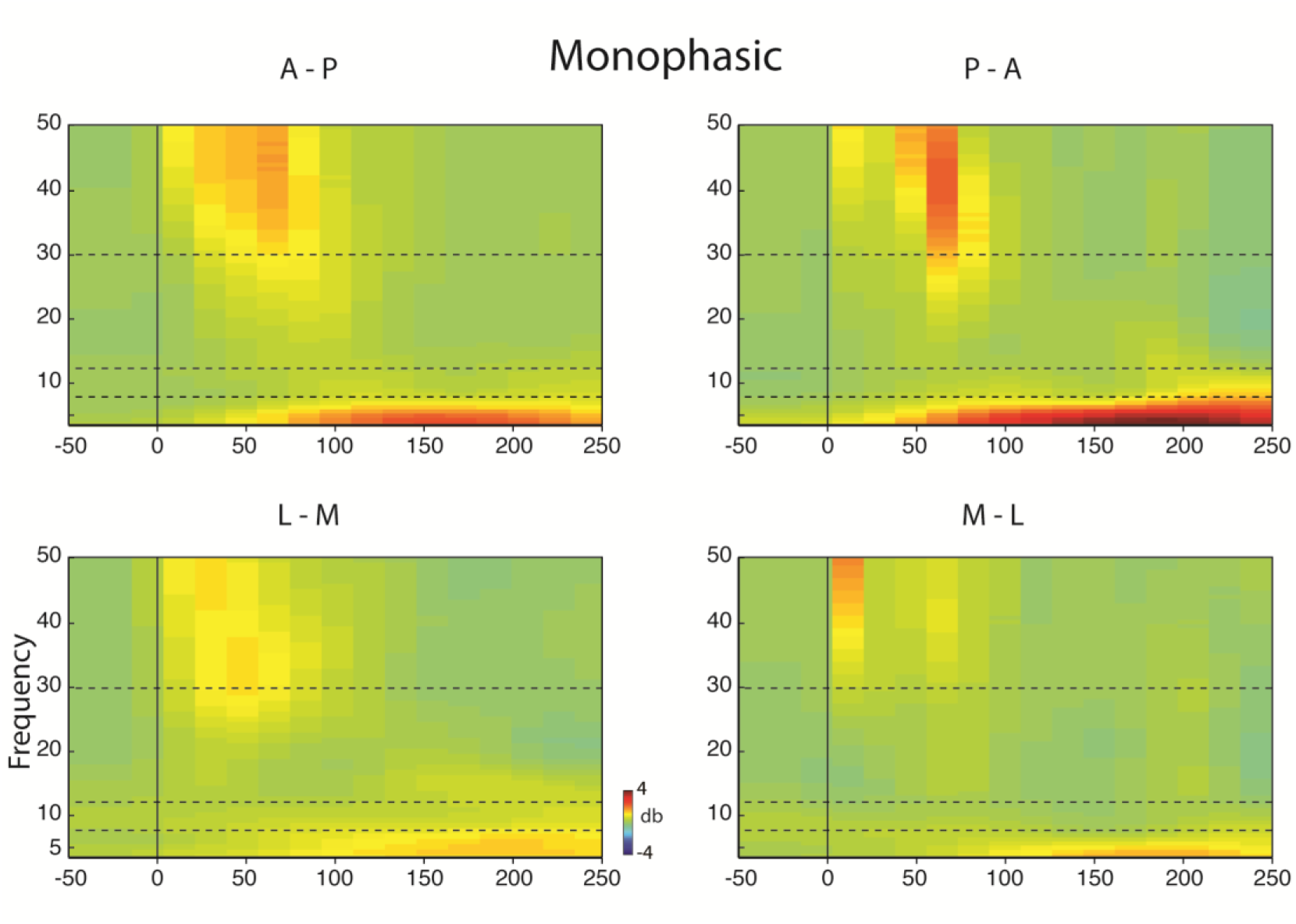
Time-frequency representation in the four monophasic directions computed over the four electrodes near the TMS coil (Fz, F2, FCz, FC2). Dashed lines represent the frequency boundaries of the 4 analyzed bands.

### Effects of coil orientation on TEPs evoked by biphasic pulses

#### Amplitude at the scalp level

Fig. 6 shows the unrectified grand average of TEPs in the four biphasic conditions. For this pulse waveform, responses in the two orientations differ in the very early components, lasting from 15 ms to 70 ms post-TMS onset. In this case, the parallel orientations, in both the L – M and M – L directions, elicits greater TEPs components. The cluster-based analysis confirmed that by delivering biphasic stimulation parallel to the gyrus, the earlier components resulted greater as compared to the perpendicular one. In particular, the M – L direction triggered a greater component peaking around 15 ms post-TMS compared to both A – P (significant cluster from 10 to 20ms, p=.01) and P – A conditions (significant cluster from 10 to 30 ms, p=.01). Moreover, the M – L component arising around 40ms post-TMS was greater than the same component in the A – P condition (significant cluster from 30 to 50ms, p=.04). The scalp distribution of the significant differences between M – L, and P – A and A – P protocols show central frontal greater amplitude for the parallel compared to the two perpendicular directions. Similarly, the L – M direction induced a greater early TEP component than the perpendicular biphasic condition in both the A – P (significant cluster from 10 to 70 ms, p<.001) and P – A (significant cluster from 10 to 20 ms, p=.02; from 30 to 60 ms, p=.016) directions. Also, these differences are distributed around frontocentral electrodes (See Fig. 7). Moreover, L – M condition elicited a greater late component when compared to the perpendicular P – A (significant cluster from 180 to 210ms, p=.009) and parallel M – L (significant cluster from 150 to 180ms, p=.028) protocols. This difference between the parallel orientations with biphasic pulses has a topography encompassing frontal and parietal left electrodes, thus contralateral with respect to the TMS hotspot (see Fig.7).

**Fig. 6:**
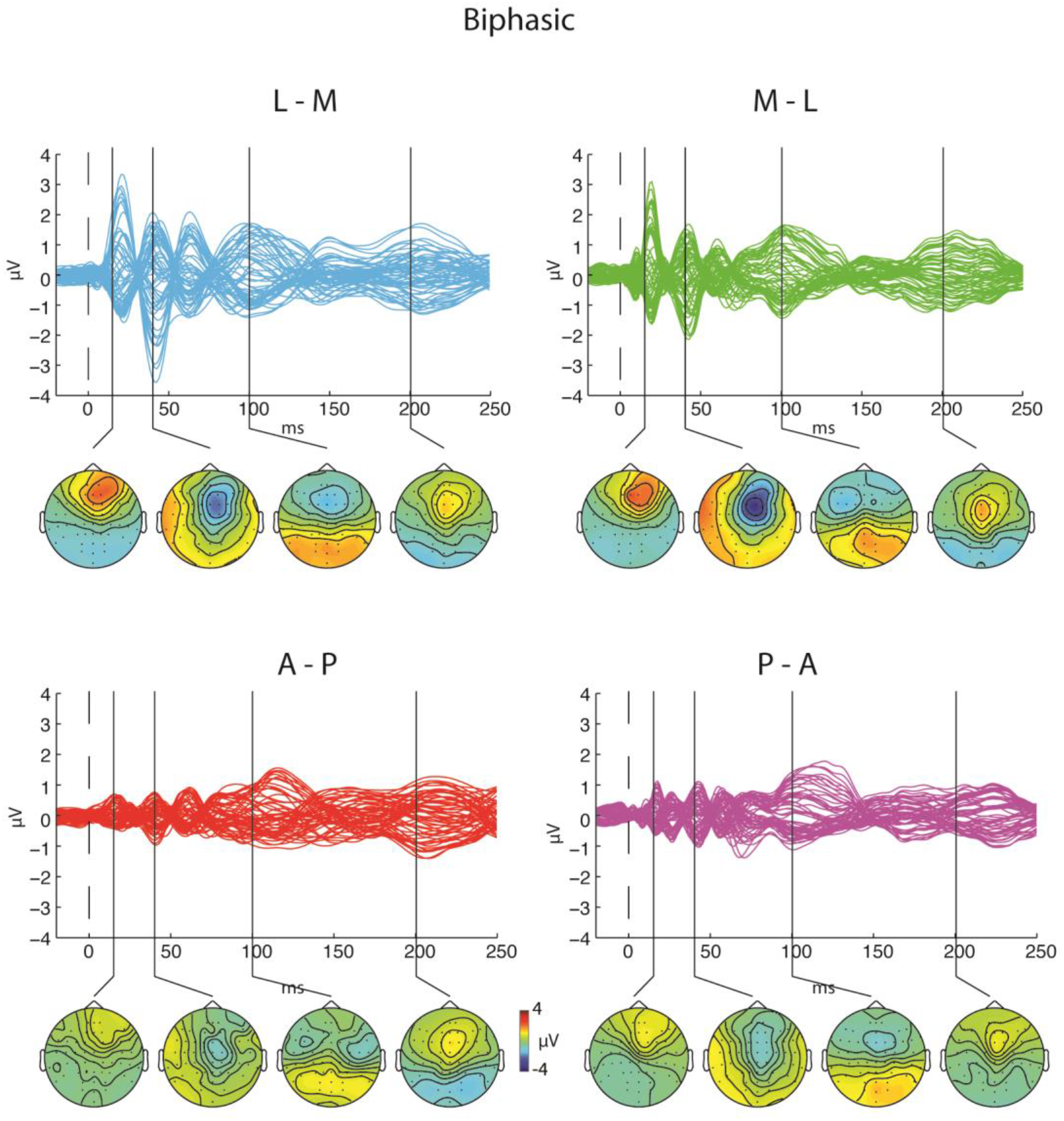
Butterfly plots and scalp topographies of TEPs grand average triggered by biphasic TMS. Top line shows results for the L – M and M – L directions while bottom line shows results for the A – P and P – A directions.

**Fig. 7:**
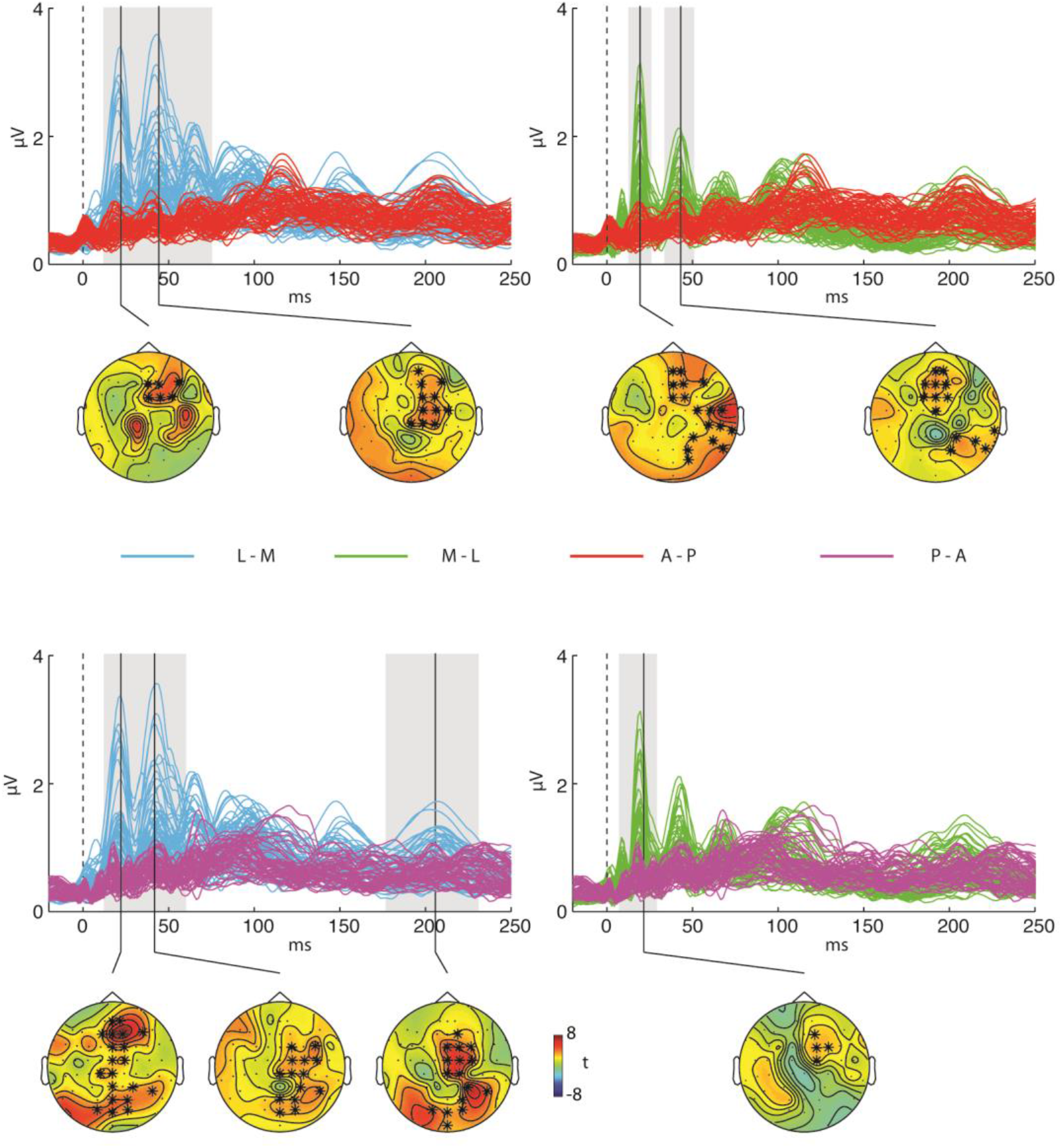
Results of the cluster based analysis for the biphasic pulse waveform. Top line displays a superimposition of rectified TEPs for the L – M and A – P (left) and M – L and A – P (right) conditions. Grey shaded areas represent time windows in which the two waveforms statistically differ. Scalp topographies represent the distribution of the statistics for each comparison with significant electrodes plotted in bold. The same is reported in the bottom line for the comparison between P – A and L – M (left) and M-L (right) protocols.

#### Cortical Source modeling

Global SCD was greater for the early time window in both parallel directions compared to the perpendicular ones (L – M vs: A – P χ^2^ (1)= 11.7; p<.001; P – A χ^2^ (1)= 15.8; p<.001; M – L vs: A – P χ^2^ (1)= 36; p=.05; P – A χ^2^ (1)= 6.8; p=.009). This greater induced cortical current in the parallel orientation was still present in the second time window (30-80ms) for the L – M direction compared to both A – P (χ^2^ (1)=5.1; p=.024) and P – A (χ^2^ (1)= 6.2; p = .012) directions, and for the M – L condition compared to the P – A direction (χ^2^ (1)=14.3; p<.001). Greater late activity was also present for the L – M stimulation compared to the A – P one (χ^2^ (1)=6.2; p=.013). No difference in SCD was present within the two parallel or perpendicular orientations. In the cortical region near the TMS target, namely right BA6, local SCD followed a similar pattern to global SCD, yet reduced in time-course. In particular, in the first time-window, L – M orientation showed a greater induced current compared to the A – P (χ^2^ (1)=11.6; p<.001) and the P – A (χ^2^ (1)=13.2; p<.001) protocols, as did the M – L condition (A – P χ^2^ (1)= 3.8; p=.05; P – A χ^2^ (1)=5.04; p=.025). In the second time-window, instead, only the L – M protocol induced greater activity compared to the A – P condition (χ^2^ (1)=5.2; p=.022). No other difference resulted significant (See Fig. 8).

**Fig. 8:**
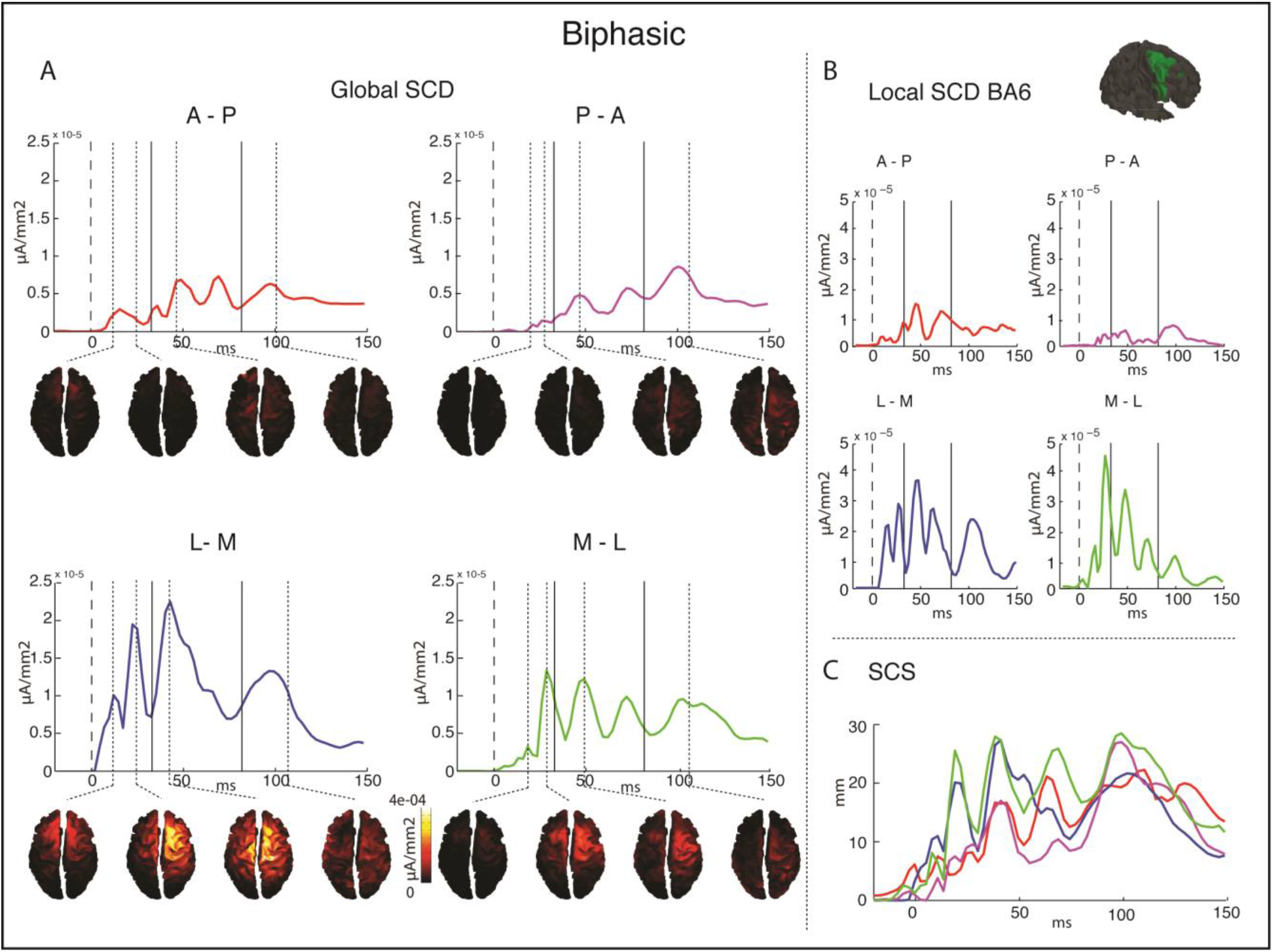
Results of the source analyses for the biphasic protocols. A) plot of the global SCD for the four directions with cortical source reconstruction at the local maxima for each of the three analyzed time windows (0-30ms; 30-80ms; 80-150ms). B) Plot of local SCD in the right BA6. C) plot of the SCS for the 4 protocols.

SCS computation showed that in the first time window, current was more widespread in the L – M condition compared to both the A – P (χ^2^ (1)=4.36; p=.037) and P – A (χ^2^ (1)=7; p=.008) protocols. Similarly, the M – L direction induced more spread cortical activity than the A – P (χ^2^ (1)= 5.8; p=.015) and P – A (χ^2^ (1)=12.5; p<.001) protocols.

#### Time-Frequency analysis

Time-Frequency analysis showed that L – M biphasic protocol resulted in greater oscillatory activity in the alpha, beta, and high beta bands compared to both A – P (α: p=.009; β: p<.001; high β: p<.001) and P – A (α: p=.005; β: p<.001; high β: p<.001) perpendicular directions. Furthermore, L – M beta activity was higher compared to the other M – L parallel condition (p=.014). M – L activity was marginally greater in the high beta band compared to the A – P direction (p=.057) (see Fig. 9).

**Fig 9.**
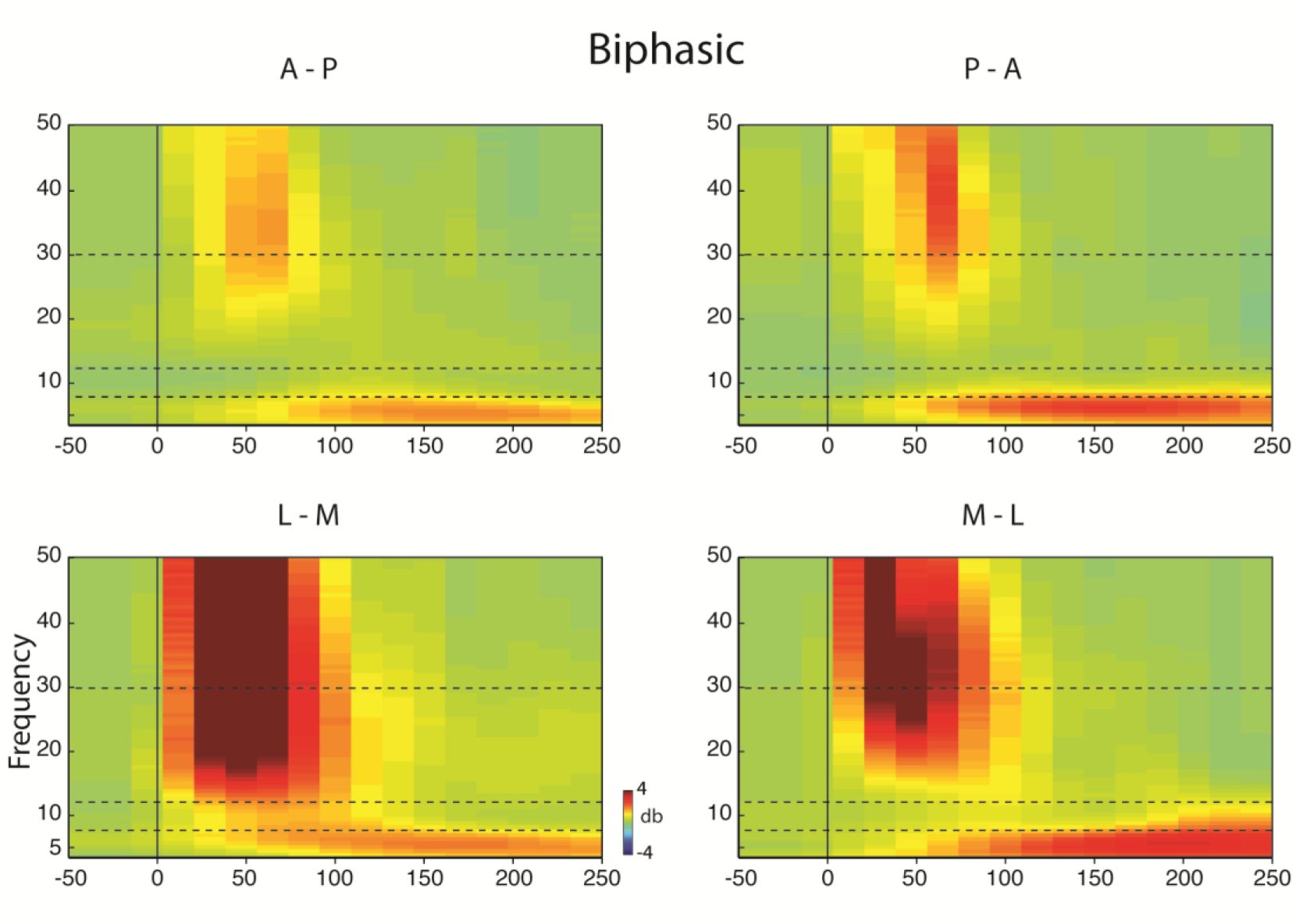
Time-frequency representation in the four biphasic directions computed over the four electrodes near the TMS coil (Fz, F2, FCz, FC2). Dashed lines represent the frequency boundaries of the 4 analyzed bands.

## Discussion

In this study, we aimed at exploring possible differences in cortical responses elicited by applying TMS over the premotor cortex with different coil orientations and pulse waveforms. To do so, we measured, using an integrated TMS-EEG system, the cortical response evoked by monophasic and biphasic TMS pulses applied with the TMS coil oriented perpendicular or parallel over the right premotor cortex, following the A – P, P – A, M – L and L – M directions. Overall, TEPs recorded after the biphasic stimulation had greater amplitude than the ones triggered by monophasic pulses. Concerning differences due to coil orientation, an analysis of TEPs time-course showed that this parameter modulated different EEG components, according to which waveform was applied. Biphasic pulses oriented L – M and M – L, i.e., parallel to the stimulated premotor gyrus, evoked a greater early component than A – P or P – A orientations, a difference which was recorded underneath the stimulation site and that was even detectable at parietal sites. Source modelling confirmed this observation, with parallel biphasic stimulation eliciting greater early cortical activity, with more widespread signal scattering, when compared to the perpendicular directions. TEPs spectral features are the typical time-frequency response evoked when stimulating the prefrontal cortex (e.g., Pellicciari et al., 2017; Rosanova et al., 2009), with an early activity peak in the beta and high beta band and a late response in the theta range. Differences in amplitude between the different directions are mirrored by an increase in the power of specific bands. Biphasic TMS applied in the M – L direction induced a diffuse greater activity in the higher frequency bands compared to perpendicular protocols and the other parallel direction.

Monophasic pulses, instead, evoked a greater middle-latency component, peaking approximately around 100 ms after TMS onset, over the stimulated area and its contralateral homolog, when oriented A – P or P – A, i.e., perpendicularly to the stimulated gyrus, than when L – M or M – L oriented. Also, in this case, source modeling confirmed greater cortical activity around 100 ms, which spread more over the cortex for the perpendicular compared to the parallel directions. Time-frequency analysis showed a general increase in all frequency bands for the A – P direction compared to the other protocols, and for the P – A compared to the M – L direction.

In general, analyses on TEP components amplitude confirmed that biphasic compared to monophasic pulses could elicit greater cortical activity at lower stimulator output levels, in line with previous reports on MEPs. Biphasic pulses, indeed, are more effective compared to monophasic ones in eliciting compound muscle action potentials (CMAPs, Niehaus et al., 2000) for both cortex and nerve stimulation, yielding a lower motor threshold, a shorter MEP latency, a steeper input/output MEP curve (Sommer et al., 2006, 2013; Sommer & Paulus, 2008; Stephani et al., 2016) and having a more complex pattern of cortical activation (Di Lazzaro et al., 2008). This greater cortical activity may be due to the broader stimulation induced by this waveform, which has been supposed to depolarize a greater neural population as compared with monophasic pulses (Arai et al., 2007), especially in its second and third cycle of the waveform (Sommer et al., 2013). Critically, by analyzing the cortical components of the elicited TEPs, we can provide direct evidence of the neurophysiological underpinning of this greater effectiveness of biphasic waveform. In the L – M and M – L stimulation direction, indeed, this type of waveform elicited a greater amplitude in the first TEP components. Early cortical EEG response to TMS has often been associated to several cortical excitability properties of the stimulated area, especially concerning M1 (Mäki & Ilmoniemi, 2010; Ziemann et al., 2015) and prefrontal stimulation (Ilmoniemi & Kičić, 2010). In particular, over the motor cortex three components have usually been reported before 100ms post TMS, peaking around 30, 45 and 60ms (Ziemann et al., 2015), and similar components have been described for the premotor cortex, even if some differences in amplitude and latency are present (Casarotto et al., 2010; Lioumis et al., 2009). The neurophysiological mechanisms generating these components have been debated in the literature. Early TEP components were modulated by tDCS when measured both from M1 (Pellicciari et al., 2013) and the prefrontal cortex (Pisoni et al., 2017), clearly building a bridge between the amplitude of these components and cortical excitability of the stimulated area. Similarly, Komssi et al. (2004) showed that these components are the most influenced by TMS intensity. Pharmacological research, instead, showed that GABAa receptors inhibition modulates an early component peaking at 45ms (Premoli et al., 2014). It has to be noted, however, that while this modulation in cortical inhibition showed a topography centered over the cortical regions contralateral to the TMS target (Premoli et al., 2014), modulation of cortical excitability effects remained under the stimulated area (Pellicciari et al., 2013; Source reconstruction in Pisoni et al., 2017). Thus, it is possible that these early components reflect different neurophysiological processes, which are related to intracortical inhibition (in regions connected to the target area) and cortical excitability of the stimulated cortical site. The present results show an increase in the early TEP components with the L-M biphasic waveform orientation which is centered over the stimulated region, supporting the hypothesis that this protocol is the most efficient in triggering action potential from pyramidal neurons directly under the stimulator, rather than a cortico-cortical response (Ilmoniemi & Kičić, 2010). In line with this, MEP research reported that L – M currents had been shown to induce D-waves, which are a reflection of direct layer V pyramidal neurons depolarization, even at low intensities (Di Lazzaro et al., 1998; for a review see Di Lazzaro et al., 2008; Di Lazzaro & Ziemann, 2013). A recruitment of this neural population might end in a greater TEP first component.

Conversely, for the monophasic pulse, the A – P and P – A directions, which run perpendicular to the targeted gyrus, were more efficient than the L-M orientation to elicit a 100 ms latency component. This component is one of the most commonly reported and more reproducible TEP component in literature (Bender et al., 2005; Bonnard et al., 2009; Kähkönen & Wilenius, 2007; Kičić, 2009; Lioumis et al., 2009; Massimini et al., 2005; Nikulin et al., 2003; Paus et al., 2001), and has been consistently associated with cortico-cortical inhibitory activity (Bikmullina et al., 2009; Premoli et al., 2014; for a review see Ilmoniemi & Kičić, 2010). Specifically, N100 amplitude has been found to be negatively associated with MEP amplitude (Kičić et al., 2008), it is modulated by coil orientation (Bonato et al., 2006) and linked with short intra-cortical inhibition (SICI) and with long intra-cortical inhibition (LICI) induced by paired-pulse TMS (Daskalakis et al., 2008; Ferreri et al., 2011; Fitzgerald et al., 2009; Rogasch & Fitzgerald, 2013). Moreover, if M1 is active while measuring TEPs or is prepared to perform an action, N100 results reduced (Bender et al., 2005; Nikulin et al., 2003), but increased when the action is inhibited (Bonnard et al., 2009). Similarly, studies applying theta burst stimulation on the prefrontal cortex showed a modulation of the 100ms component which was linked to markers of cortical inhibition as LICI (Chung et al., 2017). Pharmacological studies finally proved the link between GABAb receptors and N100 amplitude, showing that the administration of baclofen, a GABAb receptor agonist, significantly enhanced N100 amplitude (Premoli et al., 2014). Crucially, Premoli and colleagues (2014), showed that the intracortical inhibitory component linked to GABAb receptors showed a topography encompassing cortical areas contralateral to the TMS target. Similarly, we highlighted an increase in the 100ms component, which shows a scalp distribution including the stimulation site and its contralateral counterpart, suggesting greater recruitment of trans-callosal inhibitory connections running between homolog regions of the two hemispheres. Monophasic pulse perpendicularly applied with respect to the stimulated cortical gyrus, thus, seem to be the best-suited protocols to highlight, and possibly modulate, neurophysiological processes relying on such connections. In this sense, our data fit well with rTMS findings which report monophasic pulses as being more efficient in inducing inhibitory after-effects following low-frequency rTMS protocols (Sommer et al., 2002; Taylor & Loo, 2007), especially with the A – P pulse direction (Tings et al., 2005).

Finally, it has to be noted that by using the TMS-EEG approach we were able to reveal neurophysiological differences in stimulation protocols which up to now were derived from indirect measures of cortical response, such as MEP or spinal recordings and limited to M1 or V1 stimulation. This opens a whole new scenario in which effects of different stimulation protocols can be directly tested by applying TMS over the target cortical region of interest.

As a potential limitation, we have to point out that monophasic and biphasic stimulation elicit different auditory and somatosensory feelings. Even if we did not directly compare the two pulses, and applied efficient masking techniques (Ter Braack et al., 2015), we cannot exclude some influence of these effects on our results. However, as Ter Braak and collaborators (2015) demonstrated, both auditory and bone conduction-related feelings do not affect early TEP components, while they do affect the N100 complex. However, since we had a modulation of this component by tilting the monophasic pulse, while no modulation occurred with the biphasic stimulation, we feel like it is unlikely that haptic sensations are involved in this effect.

From a theoretical point of view, our results support with direct evidence the hypothesis that different pulse waveforms and directions do have different stimulation outcomes in terms of targeted neural population and induced cortical mechanisms: results showed that biphasic pulse oriented parallel to the stimulated premotor gyrus triggered cortical excitability mechanisms within the target area, while monophasic stimulation perpendicularly oriented to the target gyrus mainly triggered inhibitory interneurons. These findings may guide scientists and clinicians using TMS as their research and treatment method in designing more efficient stimulation protocols to address their scientific goals.

